# Eliminating fibroblast activation protein-α expressing cells by photoimmunotheranostics

**DOI:** 10.1101/2021.11.18.469110

**Authors:** Jiefu Jin, James D. Barnett, Balaji Krishnamachary, Yelena Mironchik, Catherine K. Luo, Hisataka Kobayashi, Zaver M. Bhujwalla

## Abstract

Photoimmunotherapy (PIT) using an antibody conjugated to a near infrared dye IR700 is achieving significant success in target-specific elimination of cells. Fibroblast activation protein alpha (FAP-α) is an important target in cancer because of its expression by cancer associated fibroblasts (CAFs) as well as by some cancer cells. CAFs that express FAP-α have protumorigenic and immune suppressive functions. Using immunohistochemistry of human breast cancer tissue microarrays, we identified an increase of FAP-α+ CAFs in invasive breast cancer tissue compared to adjacent normal tissue. We found FAP-α expression increased in fibroblasts co-cultured with cancer cells. In proof-of-principle studies, we engineered human FAP-α overexpressing MDA-MB-231 and HT-1080 cancer cells and murine FAP-α overexpressing NIH-3T3 fibroblasts to evaluate several anti-FAP-α antibodies and selected AF3715 based on its high binding-affinity with both human and mouse FAP-α. After conjugation of AF3715 with the phthalocyanine dye IR700, the resultant antibody conjugate, FAP-α-IR700, was evaluated in cells and tumors for its specificity and effectiveness in eliminating FAP-α expressing cell populations with PIT. FAP-α-IR700-PIT resulted in effective FAP-α-specific cell killing in the engineered cancer cells and in two patient-derived CAFs in a dose-dependent manner. Following an intravenous injection, FAP-α-IR700 retention was three-fold higher than IgG-IR700 in FAP-α overexpressing tumors, and two-fold higher compared to wild-type tumors. FAP-α-IR700-PIT resulted in significant growth inhibition of tumors derived from FAP-α overexpressing human cancer cells. A reduction of endogenous FAP-α+ murine CAFs was identified at 7 days after FAP-α-IR700-PIT. FAP-α-targeted NIR-PIT presents a promising strategy to eliminate FAP-α+ CAFs.

## Introduction

Near infrared photoimmunotherapy (NIR-PIT) is an emerging targeted cancer therapy in which a water-soluble, photo-stable, phthalocyanine dye IRDye700DX (IR700), is conjugated to an antibody to target cancer cells (Mitsunaga et al., 2011) or pro-tumorigenic cells in the tumor microenvironment (TME) (K. Sato et al., 2016). The antibody conjugate specifically binds to the target on the cell membrane, causing membrane damage after NIR light exposure (Jin, Krishnamachary, Mironchik, Kobayashi, & Bhujwalla, 2016; Kazuhide Sato et al., 2018). A first-in-human phase 1/2 clinical trial of NIR-PIT using cetuximab-IR700 (RM1929) to treat inoperable recurrent head and neck cancer patients that successfully concluded in 2017 was “fast-tracked” by the FDA for a phase 3 trial (https://clinicaltrials.gov/ct2/show/NCT03769506) (Kobayashi & Choyke, 2019). In September 2020, the first drug and the laser system for human use, cetuximab-IR700 (ASP1929, Akalux™) and a 690nm laser system (BioBlade™), were conditionally approved and registered for clinical use by the Pharmaceuticals and Medical Devices Agency (PMDA) in Japan, with health insurance coverage available for recurrent head and neck squamous cell carcinoma since January 2021.

Here we evaluated the use of NIR-PIT to target FAP-α expressing cells. FAP-α, a member of the dipeptidyl peptidase (DPP) family, is expressed at low or undetectable levels in normal tissue (Niedermeyer et al., 2001), but is upregulated in CAFs in human epithelial tumors. CAFs constitute the most abundant cell population in the stroma of most solid tumors (Prakash, 2016), presenting a ubiquitous target in cancer. A few studies have also reported expression of FAP-α by cancer cells such as melanoma (Fitzgerald & Weiner, 2020). Because FAP-α expressing CAFs exert pro-tumorigenic and immunosuppressive functions (Kieffer et al., 2020), FAP-α is an attractive molecular target for cancer imaging and treatment, especially in stromal-rich desmoplastic cancers (Puré & Blomberg, 2018). FAP-α is also associated with human pathologies such as fibrosis, arthritis, atheloscerosis and autoimmune diseases (Fitzgerald & Weiner, 2020). A recent study investigated the use FAP-α photodynamic therapy in experimental arthritis (Dorst et al., 2020).

Small molecule FAP-α inhibitors have been previously developed and evaluated in clinical studies. The most commonly used FAP-α inhibitor, Val-boroPro (Talabostat), failed in a number of phase II clinical trials, with limited clinical benefit even in combination with chemotherapy (Eager, Cunningham, Senzer, Richards, et al., 2009; Eager, Cunningham, Senzer, Stephenson, et al., 2009; Narra et al., 2007). The minimal efficacy of Sibrotuzumab (BIBH1), a humanized version of the murine anti-FAP-α monoclonal antibody F19 (Scott et al., 2003), resulted in a failed early phase II clinical trial in patients with metastatic colorectal cancer (Hofheinz et al., 2003). The failure of Talabostat and Sibrotuzumab in clinical trials indicated that binding of FAP-α or blocking the enzymatic activity of FAP-α was not effective in mediating clinical benefit, identifying depletion of FAP-α+ CAFs as a better strategy. The use of an FAP-α inhibitor (FAPI-04) labeled with a therapeutic radionuclide resulted in growth control of pancreatic cancer xenografts (Watabe et al., 2020). Other systemic strategies to deplete FAP-α+ stromal cells such as DNA vaccines (Loeffler, Krüger, Niethammer, & Reisfeld, 2006; Xia et al., 2016), and adenoviral vectors (de Sostoa et al., 2019; Pang et al., 2017; Zhang & Ertl, 2016), have resulted in favorable therapeutic outcomes in preclinical tumor models. Antibody-based therapeutics, such as immunotoxins (Fang et al., 2016; Ostermann et al., 2008) and antibody fragment-based chimeric antigen receptor (CAR) engineered T-cells (Kakarla et al., 2013; Lo et al., 2015; Wang et al., 2014), used to deplete FAP-α+ cells in preclinical models, also resulted in reduced tumor growth. However, FAP-α specific CAR T cells resulted in toxicity and cachexia because FAP-α+ fibroblasts play a pivotal role in preserving tissue homeostasis in the skeletal muscle, and FAP-α is also expressed by PDGFR-α^+^, Sca-1^+^ multipotent bone marrow stromal cells (BMSCs) (Tran et al., 2013). These results identified potential problems with systemic elimination of FAP-α+ stromal cells, highlighting the importance of eliminating FAP-α+ CAFs in tumors without damaging FAP-α+ cells in normal tissues.

Here, we first investigated expression of FAP-α CAFs in human breast cancer tissue microarrays and in human mammary fibroblasts (HMFs) co-cultured with human breast cancer cells. Next, in proof-of-principle studies, we engineered human FAP-α overexpressing MDA-MB-231 and HT-1080 cancer cells and murine FAP-α overexpressing NIH-3T3 mouse fibroblasts to select a high-affinity human and mouse cross-reactive anti-FAP-α antibody, and investigated the specificity and effectiveness of the antibody-IR700 conjugate to bind to and eliminate FAP-α+ cells in culture and *in vivo*. NIR emission of IR700 has a penetration depth of ~ 1 cm in tissue allowing the detection of IR700 in tumors with noninvasive fluorescence imaging to determine the optimal timing of PIT. The *in vivo* studies demonstrated the feasibility of combining target-specific antibody binding with tumor localized NIR exposure to eliminate FAP-α+ cells in solid tumors.

## Results

### FAP-α expressing CAFs increase in breast cancer TMAs compared to normal adjacent tissue

To identify FAP-α expressing CAFs in breast cancer, human breast cancer TMAs (6 cases with 2 cores per group, n = 6) and matched adjacent breast tissue were immunostained (Fig. 1a and b). The percent area occupied by FAP-α+ CAFs was ~ 4.3%, while in adjacent breast tissue cores, the fractional area of FAP-α+ CAFs was 0.41% (Fig. 1c), confirming that numbers of FAP-α+ CAFs significantly increased in breast cancer. Co-culturing HMFs with MDA-MD-231 cells resulted in an increase of FAP-α fluorescence intensity values from 9.5 ± 5.2 to 18.0 ± 8.5, as measured by flow cytometry from two independent experiments (Fig. 1- figure supplement 1).

**Figure 1.**
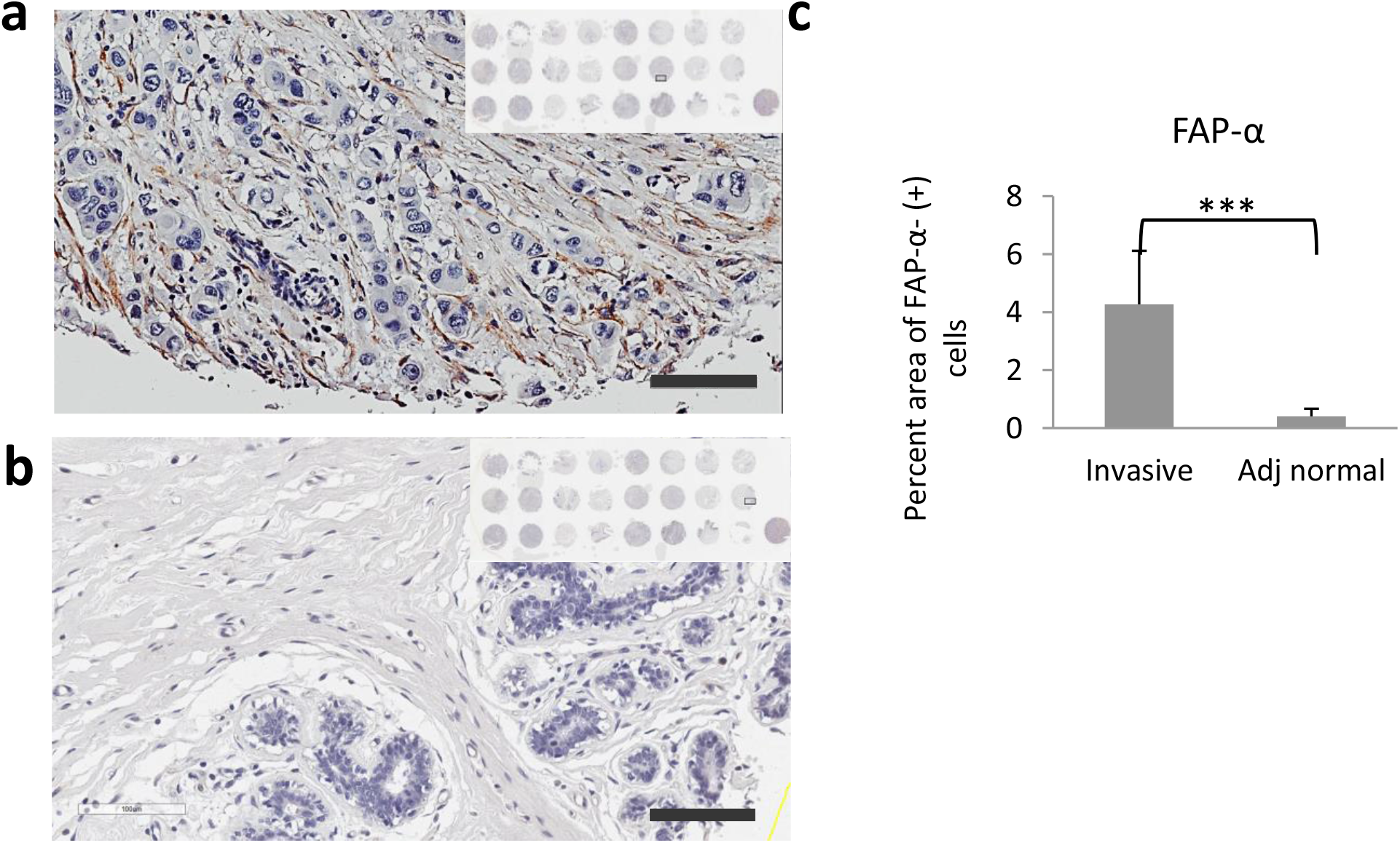
FAP-α expressing CAFs increase in breast cancer. Representative FAP-α immunostained image of breast invasive ductal carcinoma tissue (a) and the matched adjacent normal breast tissue (b) at 20X magnification, inset shows the entire TMA with rectangle box indicating the location of the image. Scale bar = 100 μm. (c) Percent area of FAP-α+ CAFs in invasive breast cancer cores compared to adjacent normal tissue cores. ***P<0.001. Values represent Mean ± SD from 6 cases with 2 cores per group (n = 6).

**Figure 1-figure supplement 1.**
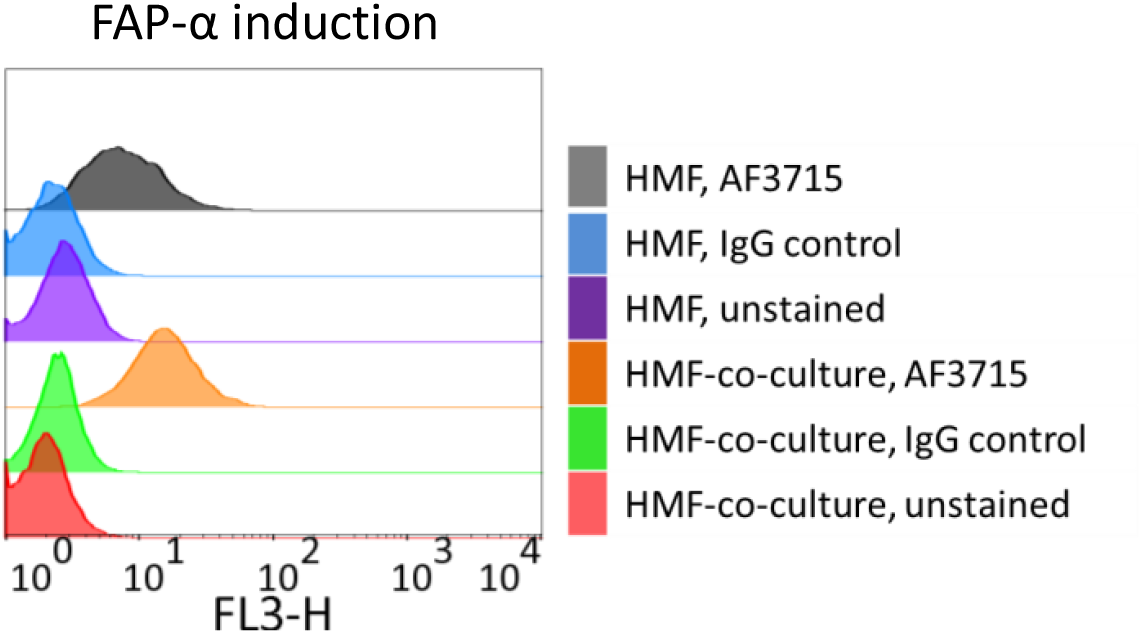
FAP-α is induced in HMFs by co-culturing with human breast cancer cells. Flow cytometry analysis confirmed induction of FAP-α in HMFs after co-culturing with 231 cells.

### Validation of FAP-α overexpressing cells and mouse and human specificity of FAP-α antibody

Increased protein expression and FAP-α mRNA were confirmed in lentivirally transduced 231-FAP, HT-1080-FAP (Fig. 2a and b) and 3T3-FAP cells (Fig. 2-figure supplement 1) compared to parent cells. Flow cytometry of live cells further confirmed increased expression of FAP-α in plasma membranes of 231-FAP and HT-1080-FAP cells, and the negligible amount of FAP-α protein on parental cell membranes (Fig. 2c). We additionally evaluated FAP-α expression by flow cytometry in two primary CAFs from pancreatic cancer (CAF35) and prostate cancer (PCAFs) (Fig. 2d), and also measured FAP-α mRNA levels in these cells (Fig. 2b). Expression levels of FAP-α determined by flow cytometry in CAF35 and PCAFs were approximately a third of those observed in the engineered cells (Fig. 2d).

**Figure 2.**
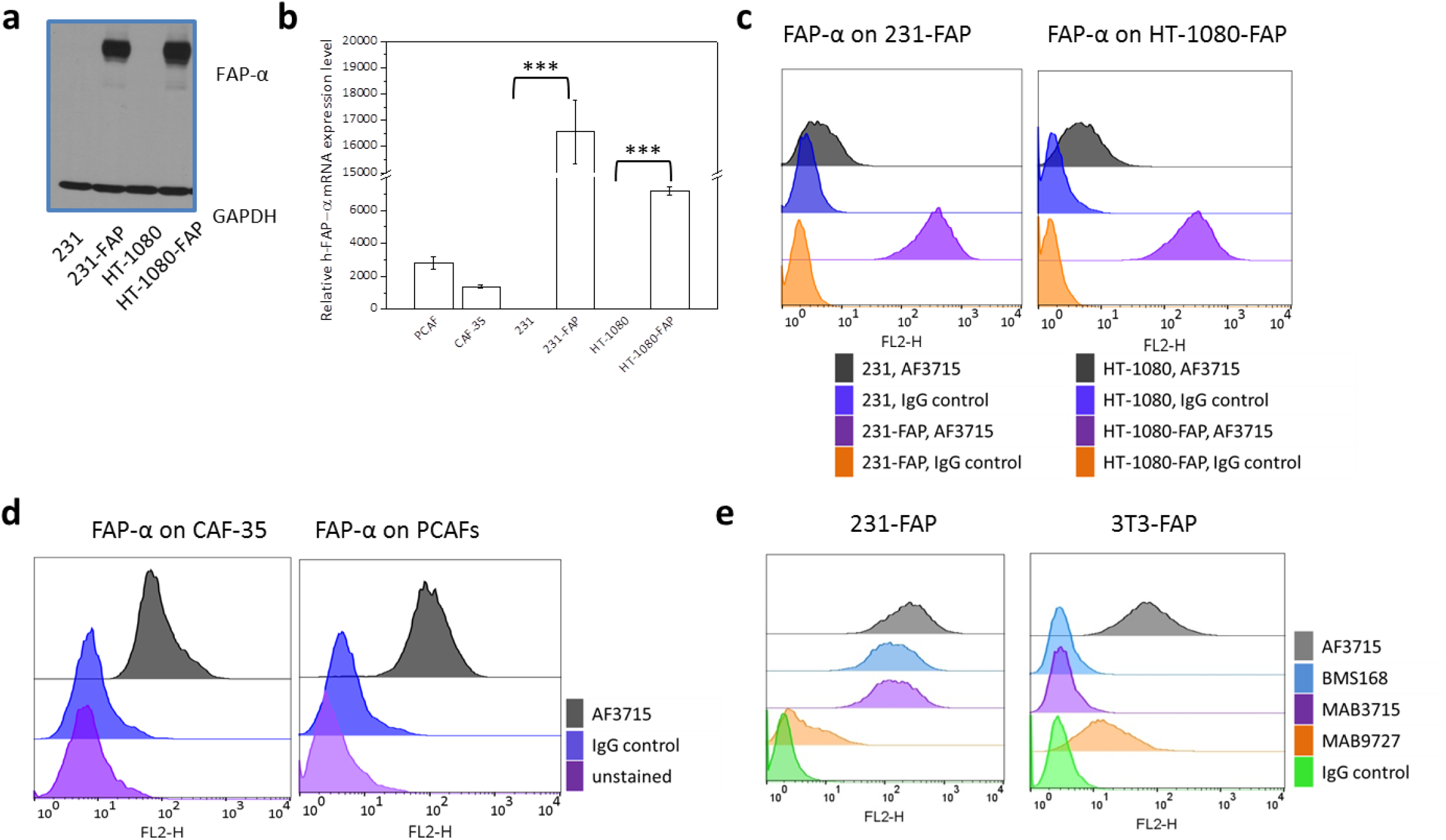
Verification of FAP-α overexpression, and human and murine FAP-α cross-reactivity of AF3715. (a) Immunoblot analysis confirming FAP-α overexpression in 231-FAP and HT-1080-FAP cells. (b) Relative FAP-α mRNA expression levels (Mean ± SD) of prostate cancer associated fibroblasts (PCAFs), pancreatic cancer associated fibroblasts (CAF35), MDA-MB-231 (231), MDA-MB-231 (231-FAP), HT-1080 and HT-1080-FAP cells. ***P<0.001. Flow cytometry analysis confirming FAP-α overexpression in 231-FAP and HT-1080-FAP cells with 231 and HT-1080 cells used as controls (c) and CAF35 and PCAFs (d). (e) AF3715 binds human and murine FAP-α with high affinity as shown by flow cytometry analysis of the binding of several anti-FAP-α antibodies to 231-FAP and 3T3-FAP. AF3715 was compared with BMS168, MAB3715, and MAB9727. Mouse IgG isotype antibody was used as control.

**Figure 2-figure supplement 1.**
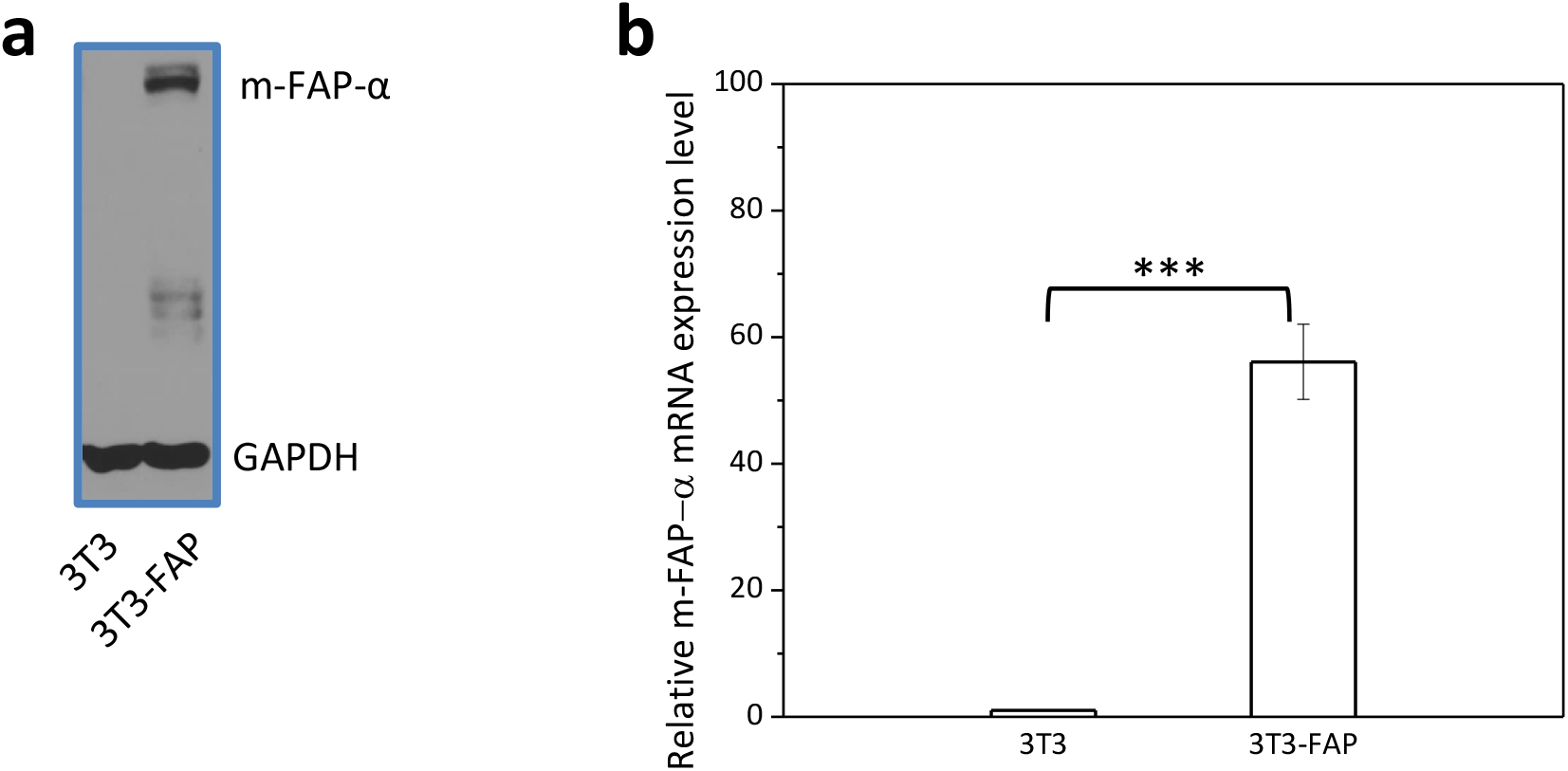
Verification of FAP-α overexpression in 3T3-FAP cells. (a) Western blotting analysis confirming FAP-α overexpression in 3T3-FAP cells. (b) Relative murine FAP-α mRNA expression levels (Mean ± SD) of NIH-3T3 and 3T3-FAP cells. ***P<0.001.

Several commercially available anti-FAP-α antibodies were evaluated with flow cytometry for their binding affinity to 231-FAP and 3T3-FAP cells. To evaluate the antibodies, identical concentrations of phycoerythrin conjugated secondary antibodies were used. Antibodies ab137549, ab218164, ab28244, ab53066, ab207178, ab227703 and PA5-51057 showed undetectable binding to either 231-FAP or 3T3-FAP (data not shown). Among the antibodies that bound to either 231-FAP or 3T3-FAP, AF3715 displayed the highest binding affinity to both human and murine FAP-α (Fig. 2e).

### Binding specificity of FAP-α-IR700

We conjugated IR700 to AF3715 to obtain the FAP-α-IR700 conjugate. Based on the spectroscopy data, the molar ratio of IR700 to antibody in FAP-α-IR700 was ~ 3.5. Confocal microscopy confirmed the selective binding of FAP-α-IR700 in 231-FAP and HT-1080-FAP cells, with negligible non-specific binding of IgG-IR700, and FAP-α – IR700 binding inhibition with 5X excess of AF3715 (Fig. 3a). Compared to the unconjugated antibody, AF3715, FAP-α-IR700 demonstrated only a slightly weaker binding affinity (Fig. 3- figure supplement 1).

**Figure 3.**
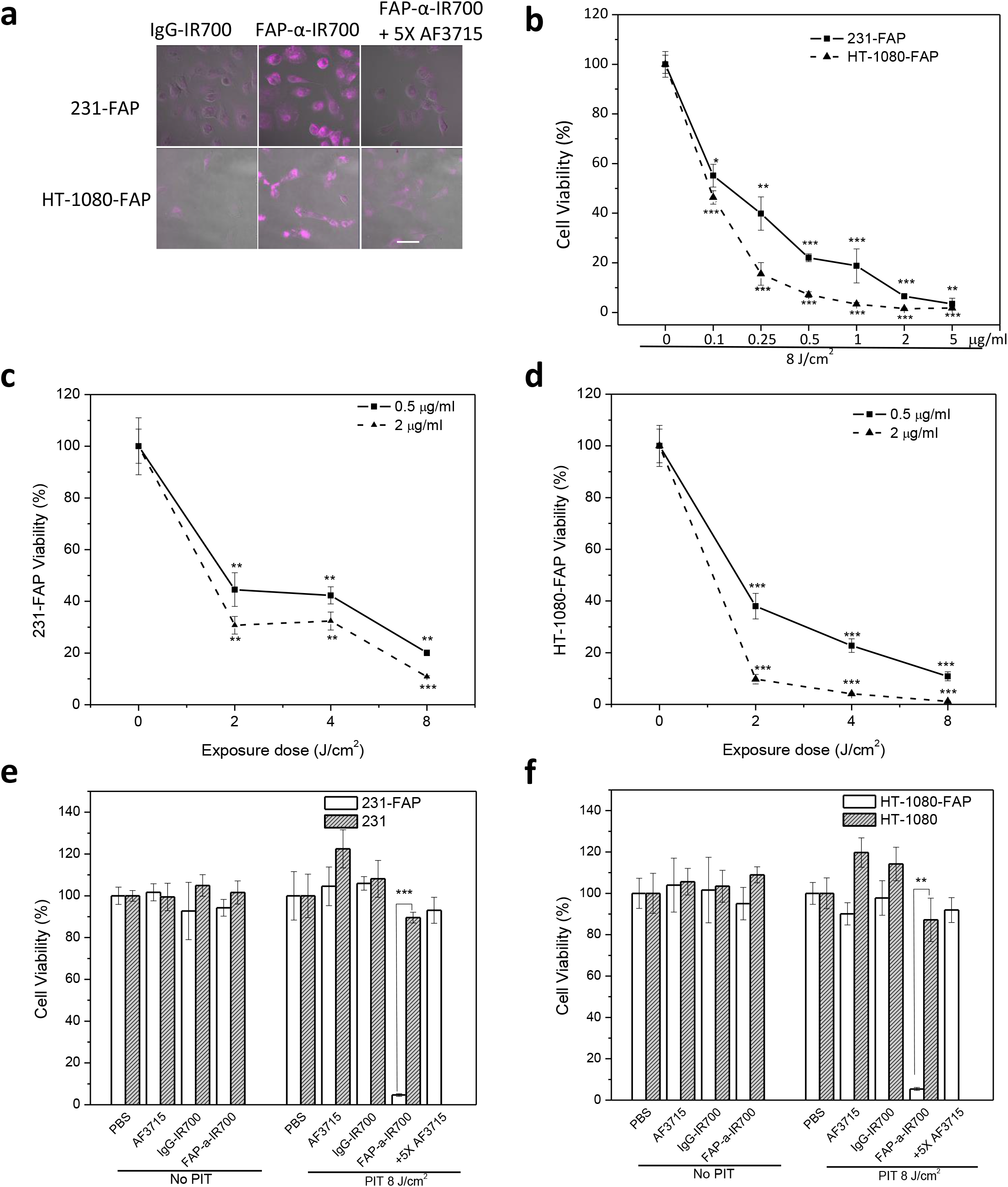
Binding specificity, dose-dependence and FAP-α-specific cell killing with FAP-α-IR700-PIT. (a) Confocal microscopy images of overlaid fluorescence and bright-field views of 231-FAP and HT-1080-FAP cells following 1-h incubation with 5 μg/ml of IgG-IR700, 5 μg/ml of FAP-α-IR700, or 5 μg/ml of FAP-α-IR700 together with 25 μg/ml of AF3715 at 37 ° C. Scale bar = 50 μm. FAP-α-IR700-mediated phototoxicity is dependent on the concentration of FAP-α-IR700 (b) and light exposure dose (c, d). FAP-α-specific cell killing only occurs with FAP-α-IR700 binding and light exposure of 231-FAP cells (e) or HT-1080-FAP cells (f). FAP-α-IR700-PIT-mediated phototoxicity is inhibited by 5X AF3715. Values represent Mean ± SD (n = 4, P<0.05, **P<0.01, ***P<0.001, for treated groups compared to PBS groups).

**Figure 3–figure supplement 1.**
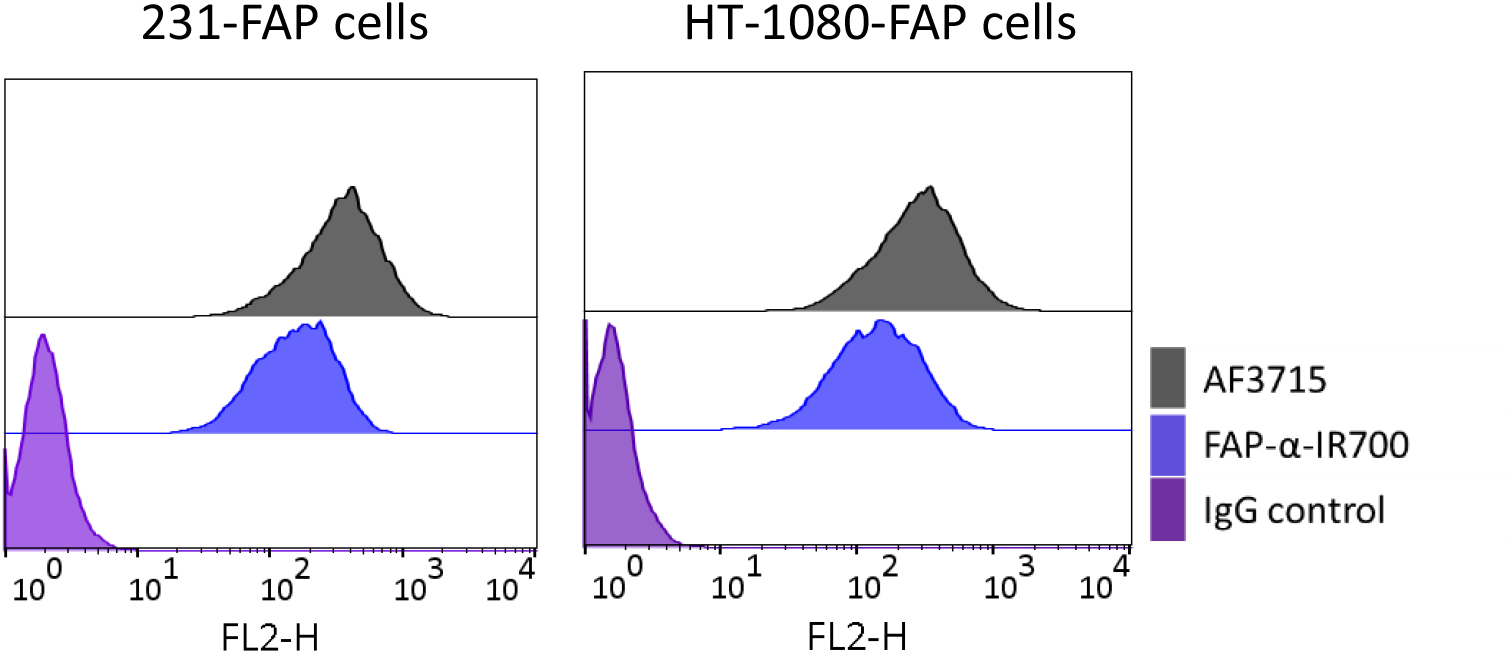
Flow cytometry analysis to determine binding of FAP-α-IR700 to 231-FAP (left) and HT-1080-FAP cells (right) when compared with AF3715 or IgG control.

### Dose-dependent and FAP-α-specific cell killing by FAP-α-IR700-PIT

Cells were incubated with FAP-α-IR700 at concentrations ranging from 0.1 to 5 μg/ml and exposed to a fixed light exposure density of 8 J/cm^2^ to confirm concentration-dependent phototoxicity in both 231-FAP and HT-1080-FAP cells (Fig. 3b). EC_50_ values of FAP-α-IR700 at 8 J/cm^2^ were ~ 0.1 μg/ml (0.67 nM) for 231-FAP and HT-1080-FAP cells (Fig. 3b). Despite comparable FAP-α expression, HT-1080-FAP cells were intrinsically more sensitive to FAP-α-IR700-PIT than 231-FAP cells. With the concentration of FAP-α-IR700 set at 0.5 or 2 μg/ml, cell death increased as light exposure intensity was increased from 2 to 8 J/cm^2^ (Fig. 3c and d). The amount of conjugate bound to the cell surface and the light exposure intensity were the two main determinants of FAP-α-IR700-PIT effectiveness. FAP-α-IR700-PIT resulted in greater than 95% cell death in both 231-FAP and HT-1080-FAP cells at 5 μg/ml and 8 J/cm^2^, and was inhibited by 5X AF3715 (Fig. 3e and f). Under dark conditions, neither AF3715 nor the conjugates showed any toxicity in the four cell lines investigated.

FAP-α-IR700-PIT also resulted in concentration and light exposure-dependent cell death in CAF35 and PCAFs, for which EC_50_ values at 8 J/cm^2^ were approximately 1 and 5 μg/ml, respectively (Fig. 4a, and b). The higher EC_50_ values in CAF35 and PCAF cells were attributed to their relatively lower FAP-α expression levels compared to 231-FAP or HT-1080-FAP cells as evident from the qRT-PCR and flow cytometry results. Similar to the cancer cells, FAP-α-specific cell killing with FAP-α-IR700-PIT was also observed in CAF35 or PCAFs and was inhibited by incubation with 5X AF3715 (Fig. 4c).

**Figure 4.**
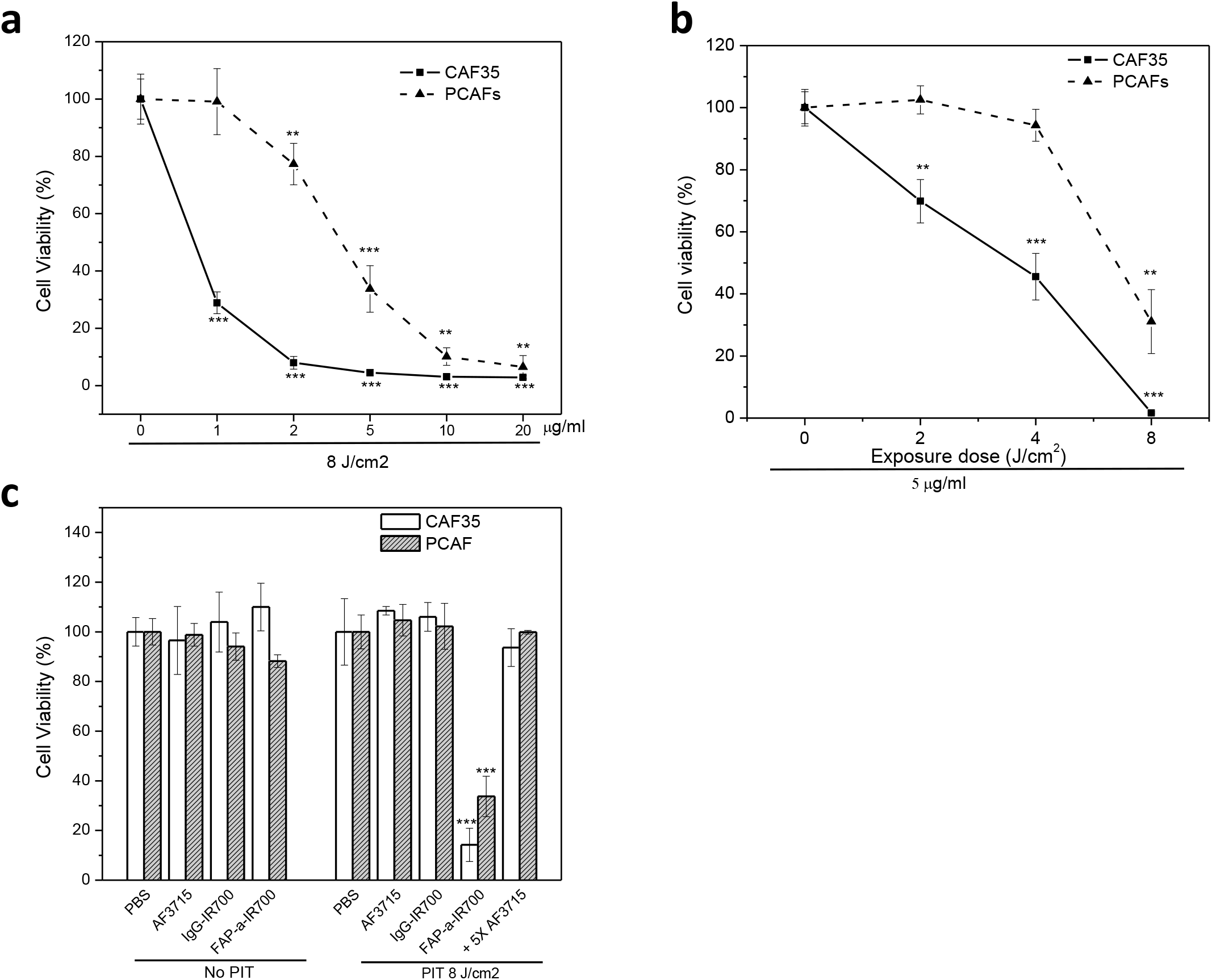
FAP-α-IR700-PIT of human CAFs. FAP-α-IR700-PIT-mediated phototoxicity of CAF35 cells and PCAFs is dependent on the concentration of FAP-α-IR700 (a) and light exposure dose (b). (c) FAP-α-specific cell killing only occurs with FAP-α-IR700 binding combined with light exposure. FAP-α-IR700-PIT-mediated phototoxicity of CAF35 and PCAFs is inhibited by incubation with 5X AF3715. Values represent Mean ± SD (n = 4, P<0.05, **P<0.01, ***P<0.001, for treated groups compared to PBS groups).

### *In vivo and ex vivo* NIR fluorescence imaging and biodistribution

At 24 h *p*.*i*., FAP-α-IR700 accumulated in 231-FAP and HT-1080-FAP tumors but not in 231 and HT-1080 tumors. Consistent with the non-specific uptake, IgG-IR700 was present at a relatively low level in all tumors (Fig. 5a and b). *Ex vivo* fluorescence images of resected tumors at 24 h *p*.*i*. confirmed the preferential accumulation of FAP-α-IR700 in 231-FAP and HT-1080-FAP tumors and the low uptake of IgG-IR700 in all the tumors (Fig. 5c and d). Quantitative uptake data, obtained from *ex vivo* fluorescence intensities normalized to the weights of organs and tumors, were presented as the percent injected dose per gram of tissue (%ID/g) (Fig. 5 e and f). At 24 h *p*.*i*., FAP-α-IR700 uptake was 1.74% and 0.84% ID/g in 231-FAP and 231 tumors, respectively (n = 4, P = 0.013); IgG-IR700 was present at 0.55% and 0.59% ID/g in 231-FAP and 231 tumors, respectively (n = 4, P = 0.65). In 231 tumors, the uptake of FAP-α-IR700 was significantly higher than the uptake IgG-IR700 (n = 4, P = 0.040) that can be attributed to the presence of endogenous murine FAP-α+ CAFs. A significantly higher uptake of FAP-α-IR700 was found in HT-1080-FAP tumors compared with HT-1080 tumors (2.23% vs 1.01% ID/g, n = 3, P = 0.025), while no difference in the uptake of IgG-IR700 was found between HT-1080-FAP and HT-1080 tumors (0.63% vs 0.56% ID/g, n = 3, P = 0.30). As with the 231 tumors, in HT-1080 tumors, the uptake of FAP-α-IR700 was significantly higher than IgG-IR700 (n = 3, P = 0.0022). In normal tissues, FAP-α-IR700 showed an uptake comparable to IgG-IR700.

**Figure 5.**
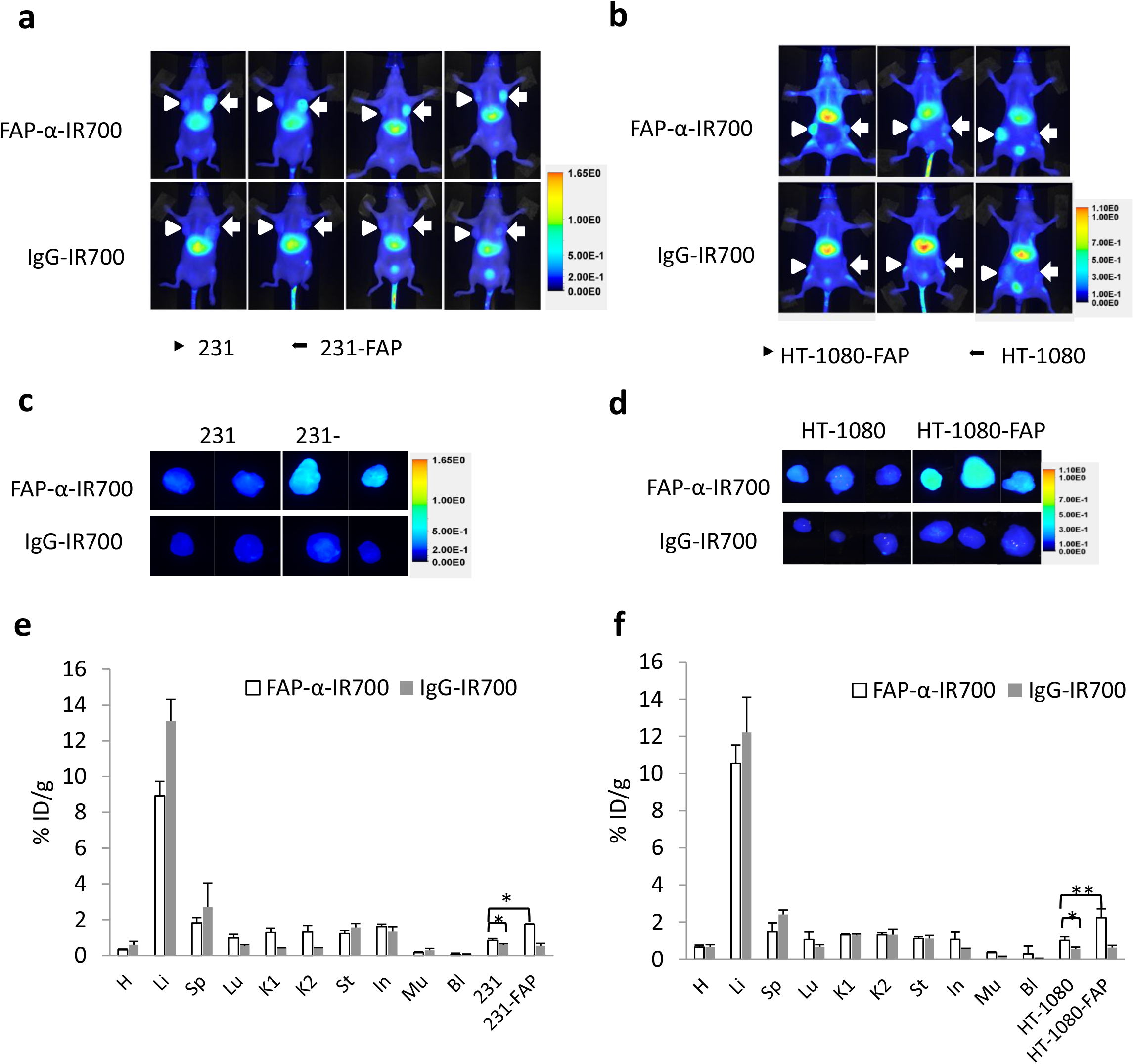
Preferential accumulation of FAP-α-IR700 in 231-FAP and HT-1080-FAP tumors. (a) NIR fluorescence *in vivo* images of mice bearing bilateral tumors (arrow head points to 231 tumor; arrow points to 231-FAP tumor) at 24 h *p*.*i*. (b) NIR fluorescence *in vivo* images of mice bearing bilateral tumors (arrow head points to HT-1080-FAP tumor; arrow points to HT-1080 tumor) at 24 h *p*.*i*. 50 μg of FAP-α-IR700 or IgG-IR700 injected *i*.*v. Ex vivo* NIR fluorescence images of resected 231 and 231-FAP tumors (c) and resected HT-1080 and HT-1080-FAP tumors (d) at 24 h *p*.*i*. Distribution of antibody conjugates in organs, 231 and 231-FAP tumors (e) and HT-1080 and HT-1080-FAP tumors (f) at 24 h *p*.*i*. Values (Mean ± SD) are normalized to % injected dose/g (%ID/g) from three or four mice per group (n = 3 or 4), *P<0.05, **P<0.01.

### *In vivo* FAP-α-IR700-PIT causes growth delay and cell death in 231-FAP and HT-1080-FAP tumors

Tumors resected from the FAP-α-IR700 group were significantly smaller than the control PBS or IgG-IR700 group at the end of the two-week treatment (Fig. 6a and b). Compared to 231-FAP tumors, HT-1080-FAP tumors responded better to FAP-α-IR700-PIT as evident from the smaller sizes, consistent with the *in vitro* NIR-PIT results (Fig. 3c and d). Tumor growth curves obtained over two weeks confirmed that FAP-α-IR700-PIT significantly inhibited the growth of 231-FAP (FAP-α-IR700 group *vs* IgG-IR700 group: at day 7, P = 0.019; at day 11, P = 0.033; at day 14, P = 0.00085) and HT-1080-FAP (FAP-α-IR700 group *vs* IgG-IR700 group: at day 7, P = 0.030; at day 11, P = 0.014; at day 14, P = 0.0063) tumors; IgG-IR700 showed no significant effect on tumor growth as compared to the PBS group (Fig. 6c and d). In 231-FAP tumors, the ratios of viable/necrotic area measured in H&E stained sections were 2.0 and 37.0 for the FAP-α-IR700 and IgG-IR700 groups (P = 0.00029), respectively (Fig. 6e). In HT-1080-FAP tumors, the ratios were 0.6 and 12.5 for the FAP-α-IR700 and IgG-IR700 group (P = 0.00048), respectively (Fig. 6f). These data confirmed that FAP-α-IR700-PIT caused significant FAP-α specific cell death in 231-FAP and HT-1080-FAP tumors.

**Figure 6.**
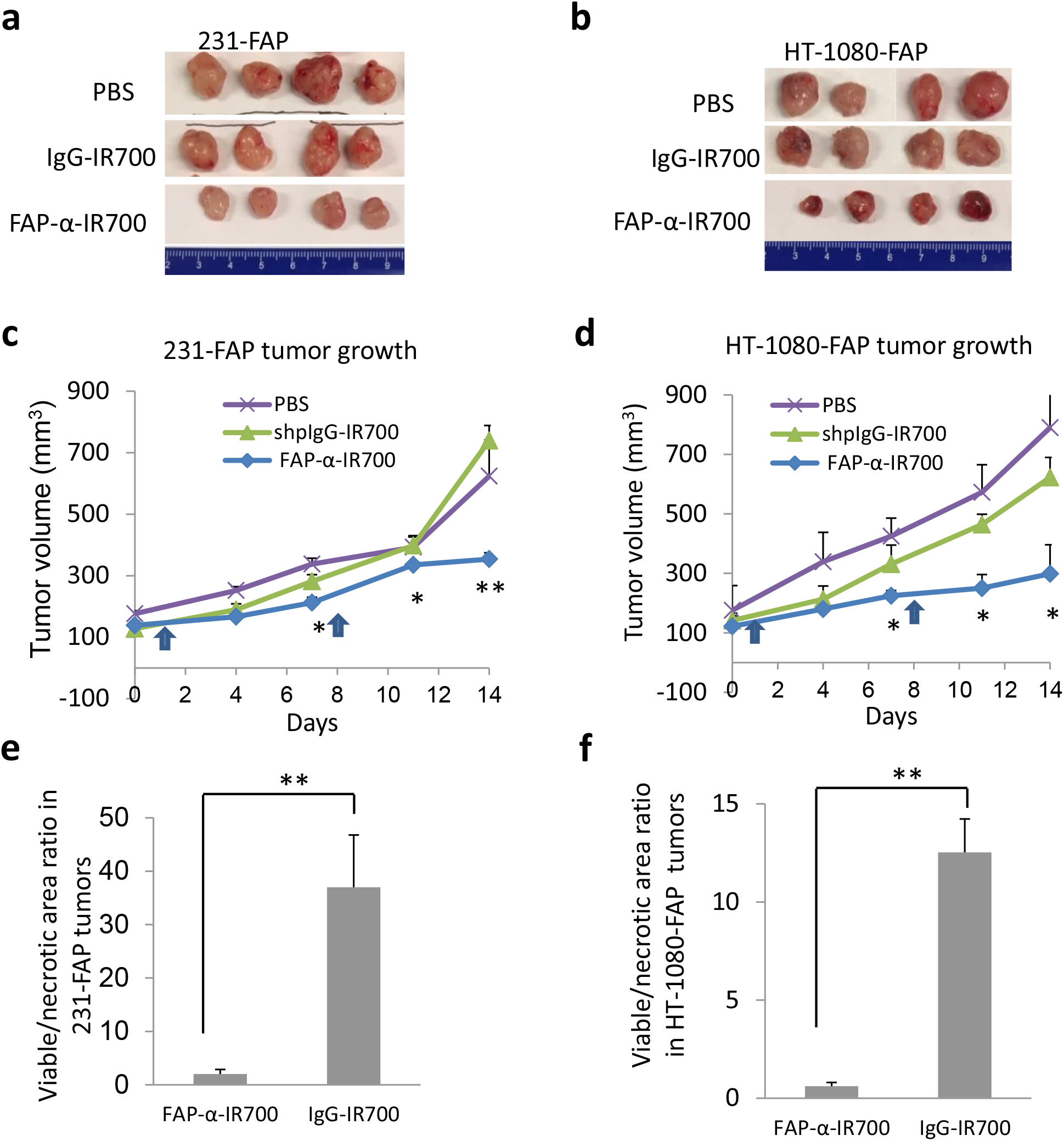
Tumor growth delay and necrosis due to FAP-α-IR700-PIT. Photographs of 231-FAP (a) and HT-1080-FAP (b) tumors resected from different groups at the end of treatment. Growth curve of 231-FAP tumors (c) and HT-1080-FAP tumors (d) in the FAP-α-IR700, IgG-IR700 and PBS groups over the duration of two weeks after injection. 100 μg of FAP-α-IR700 or IgG-IR700 or 100 μl of PBS was injected *i*.*v*. on day 0 and on day 7. Light was delivered at 200 J/cm^2^ 24 h following each injection. Values represent Mean ± SD from four mice per group. *P<0.05, **P<0.01 for the FAP-α-IR700 group compared to PBS group. The viable/necrotic area ratio in H&E stained 231-FAP (e) and HT-1080-FAP (f) tumor sections from FAP-α-IR700 and IgG-IR700 groups. Values represent Mean ± SD from four mice per group, **P<0.01 for the FAP-α-IR700 group compared to the IgG-IR700 group.

### The effect of FAP-α-IR700-PIT on FAP-α+ murine CAFs

We evaluated the effectiveness of FAP-α-IR700-PIT in depleting FAP-α+ murine CAFs in tumors *in vivo*. FAP-α-IR700-PIT was performed with tumors derived from wild-type 231 cells that do not express FAP-α. In these tumors, CAFs are of mouse origin. Compared to IgG-IR700-PIT, FAP-α-IR700-PIT resulted in a reduction of FAP-α protein in 231 tumors as identified in the western-blots (Fig. 7a). A reduction of FAP-α+ murine CAFs was also confirmed in IHC studies (Fig. 7b and c), where the percent area of FAP-α+ cells in FAP-α-IR700-PIT treated 231 tumors was significantly lower than in the IgG-IR700 group. FAP-α-IR700-PIT did not result in significant growth inhibition of 231 tumors as compared to IgG-IR700-PIT (Fig. 7-figure supplement 1), which is likely due in part to the low abundance of murine CAFs in 231 tumors, and due to their limited role in tumor growth in immune deficient mice. Flow cytometry of single cell suspensions of cells dissociated from 231 tumors revealed that 5.54% of 231 tumor-dissociated cells were FAP-α+ in contrast to 76.2% in 231-FAP tumors (Fig. 7-figure supplement 2).

**Figure 7.**
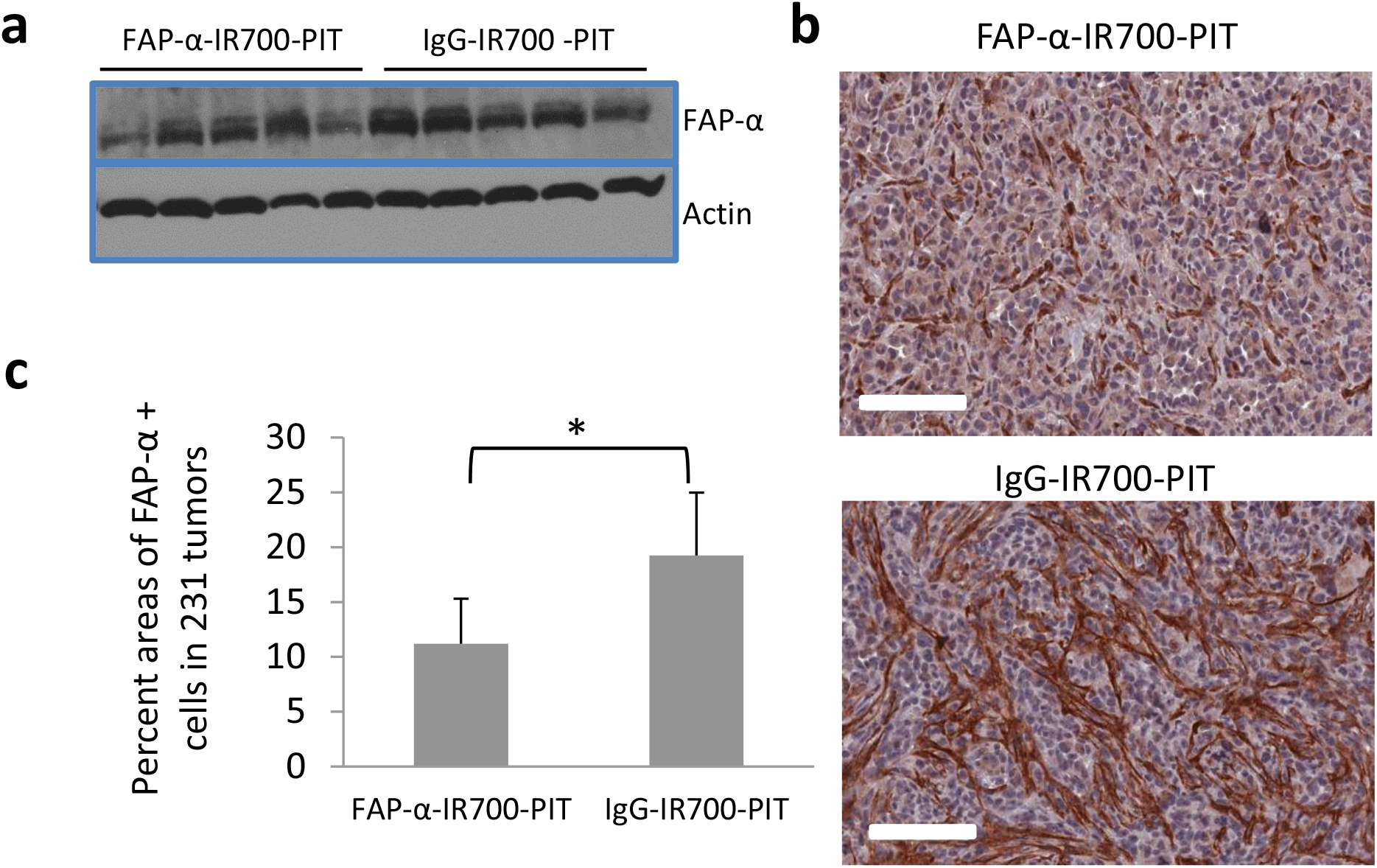
The effect of FAP-α-IR700-PIT on FAP-α+ murine fibroblasts in 231 tumors. (a) Immunoblots identify reduction of FAP-α in FAP-α-IR700-PIT treated tumors. (b) Representative FAP-α immunostained 5 μm sections from 231 tumors following FAP-α-IR700-PIT or IgG-IR700-PIT. 100 μg of FAP-α-IR700 or IgG-IR700 were injected *i*.*v*. in 231 tumor-bearing mice on day 0 and on day 7 and tumors were excised 7 days later. Light exposure was delivered at 200 J/cm^2^ 24 h after each injection. Scale bar = 100 μm. (c) Analysis of FAP-α immunostaining in 231 tumors following FAP-α-IR700-PIT or IgG-IR700-PIT. Values represent Mean ± SD from five mice per group (n = 3), *P<0.05.

**Figure 7–figure supplement 1.**
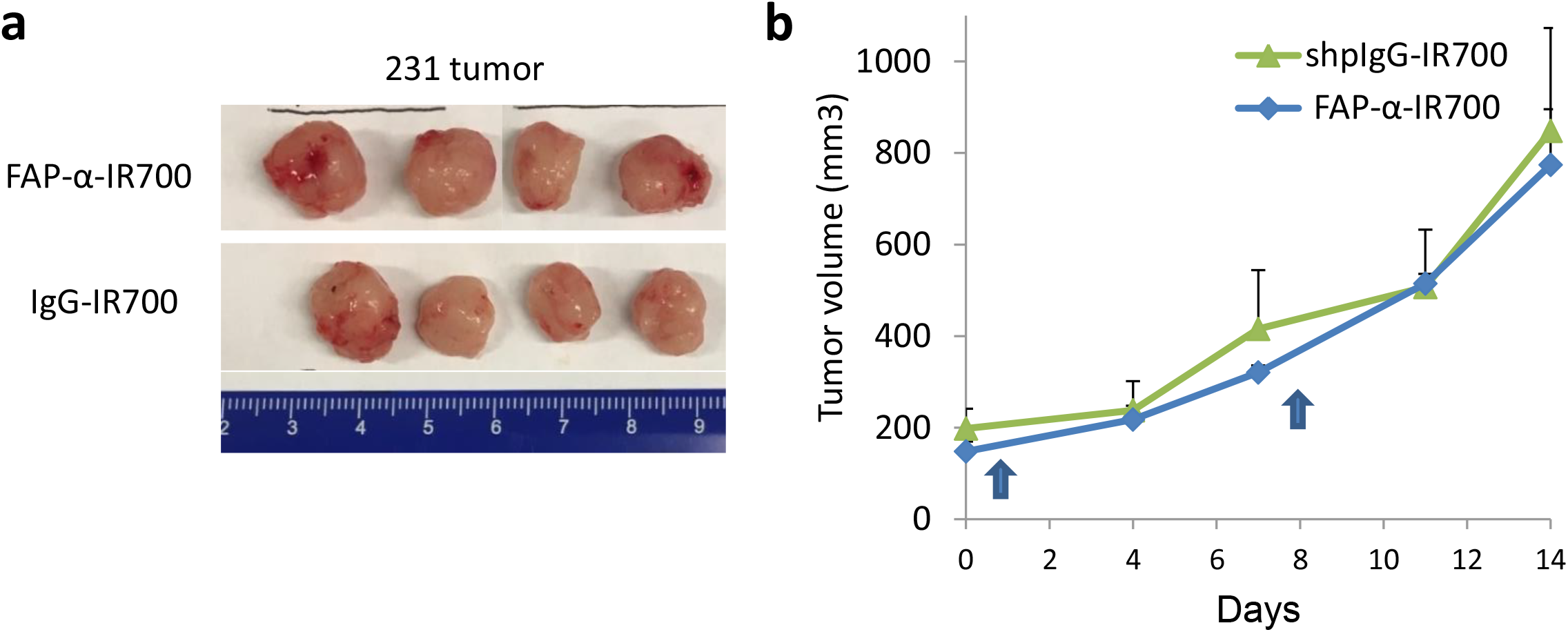
The effect of FAP-α-IR700-based PIT on 231 tumor growth. (a) Photograph of 231 tumors resected from different groups following treatment. (b) Growth curve of 231 tumors in FAP-α-IR700 and IgG-IR700 groups over the duration of two weeks. 100 μg of FAP-α-IR700 or IgG-IR700 were injected *i*.*v*. on day 0 and on day 7. Light exposure was delivered at 200 J/cm^2^ 24 h after each injection. Values represent Mean ± SD from four mice per group.

**Figure 7–figure supplement 2.**
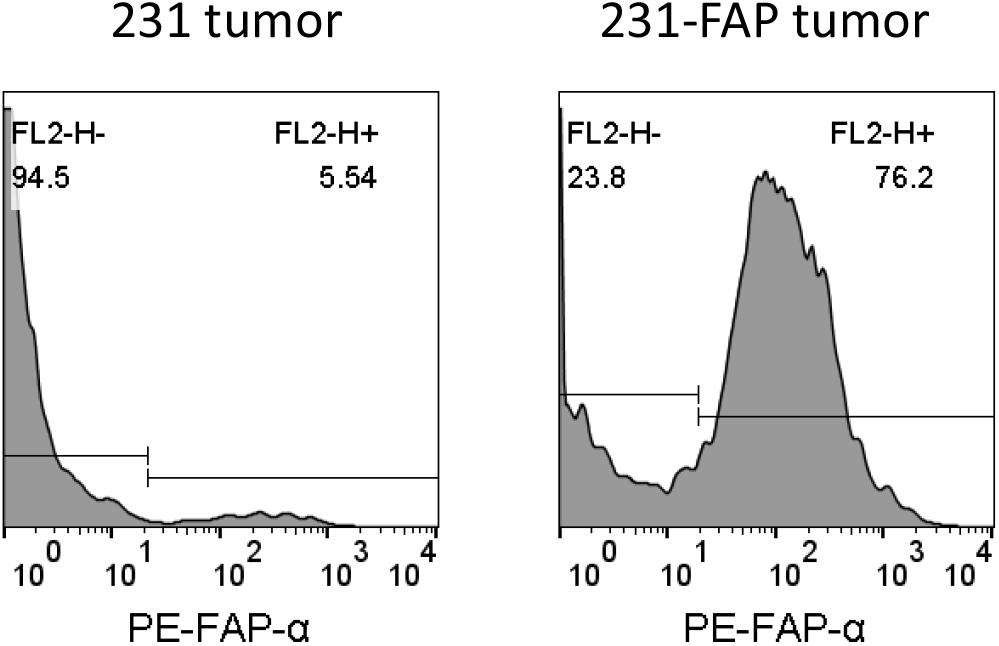
The abundance of FAP-α+ cells in 231 (left) and 231-FAP tumors (right) studied by flow cytometry.

## Discussion

We identified an increase of FAP-α+ CAFs in human breast cancer tissue compared to adjacent normal tissue, and an induction of FAP-α in HMFs co-cultured with breast cancer cells consistent with earlier studies that have detected an increase of FAP-α in breast cancer (Puré & Blomberg, 2018).

Two human cancer cell lines (MDA-MB-231 and HT-1080) and the murine fibroblast cell line, NIH-3T3, were engineered to overexpress FAP-α. Using the engineered cells, we evaluated several anti-FAP-α antibodies to ultimately select AF3715 based on its high affinity with both human and mouse FAP-α. The human and mouse cross-reactivity was important because CAFs in the mouse models are of mouse origin. To the best of our knowledge, our work is the first example of NIR-PIT using human/mouse FAP-α cross-reactive antibody.

We found that the effectiveness and specificity of NIR-PIT depended upon the number of antibody conjugates bound to the target that was mainly determined by two factors: the affinity of the antibody conjugate, and the expression level of the target at the cell surface. CAF35 and PCAFs had lower expression of FAP-α than 231-FAP and HT-1080-FAP cells in terms of mRNA and surface protein levels, and therefore required correspondingly higher concentration of FAP-α-IR700 to achieve comparable cell death. EC50 values of FAP-α-IR700-PIT in 231-FAP and HT-1080-FAP cells at 8 J/cm^2^ were ~ 0.1 μg/ml, while the values with CAF35 and PCAFs were approximately 1 and 5 μg/ml, respectively.

IR700 provided both imaging and therapeutic abilities. Detection of FAP-α-IR700 with NIR fluorescence imaging allowed noninvasive imaging of the biodistribution within the body, and detection of tumor delivery and retention, to optimally time exposure of the tumor to therapeutic light. In bilaterally inoculated wild-type and FAP-α overexpressing tumors, NIR fluorescence imaging clearly identified increased retention of FAP-α-IR700 in FAP-α overexpressing tumors compared to the corresponding wild-type tumors. By normalizing the fluorescence intensity to the weight and injected dose, we also observed increased retention of FAP-α-IR700 uptake in wild-type MDA-MB-231 and HT-1080 tumors compared to the retention of IgG-IR700, confirming the ability of FAP-α-IR700 driven fluorescence to detect endogenous FAP-α+ murine CAFs.

FAP-α-based-PIT has been previously performed with human esophageal squamous fibroblasts expressing FAP-α (Watanabe et al., 2019). Different from our studies, fibroblasts were first pre-incubated with the antibody conjugate and then inoculated together with cancer cells. NIR-PIT was given immediately following inoculation before the tumor was established. The same model system was used in a recent study to demonstrate that FAP-α-targeted NIR-PIT reduced therapeutic resistance to 5-fluorouracil in CAF co-inoculated human esophageal tumors (Katsube et al., 2021). While these previous studies provided valuable information, one study indicates that immunofluorescence from co-inoculated CAFs disappeared within 13 days, highlighting a limitation of this strategy for investigating established tumors (Fabris et al., 2010). In addition, the antibody conjugate in these studies exhibited a lower potency *in vitro*, with 20 µg/ml of the antibody conjugate at 15 J/cm^2^ resulting in a 58.9% reduction in viability of CAFs. In our study, we achieved 95% and 70% reduction in viability of CAF35 cells PCAFs, respectively with 5 µg/ml of the antibody conjugate at 8 J/cm^2^.

IHC and immunoblotting detected a decrease of endogenous FAP-α+ CAFs with FAP-α-IR700-PIT, although wild type tumor growth was not affected. This was anticipated, in part, since FAP-α+ CAFs mainly play an important role in immunosuppression (Cremasco et al., 2018; Kieffer et al., 2020; Kraman et al., 2010; Yang et al., 2016; Zhang & Ertl, 2016). Our data are consistent with an earlier study performed with FAP-α targeted nanoparticle-based phototherapy of 4T1 syngeneic tumors in immunocompetent mice (Zhen et al., 2017). In this study the suppression of tumor growth was attributed to a significant increase of CD8+ T cells. Our studies were performed with human tumor xenografts in immune suppressed mice that lack T cells.

Since FAP-α-targeted depletion by NIR-PIT has the potential to combat immunosuppression and activate systemic anti-tumor immunity in primary tumors, distant metastatic tumors not exposed to PIT may also come under immune surveillance. Future studies should evaluate FAP-α-specific NIR-PIT in syngeneic mouse models either singly or in combination with cancer immunotherapy with immune checkpoint inhibitors, to achieve effective primary and metastatic tumor control. Since CAFs actively modulate the ECM, angiogenesis, and cell migration and growth, our FAP-α-based NIR-PIT can be combined with conventional cancer cell-centric therapies to halt tumor progression and overcome drug resistance. One major limitation of PIT is that NIR light at 690 nm can penetrate and treat cancers at a depth of approximately 1 cm. Applying NIR-PIT in an intra-operative setting or by using interstitial NIR light delivered through fiber-optic diffusers inside catheter needles (Okuyama et al., 2018) or endoscopes (Nagaya et al., 2018), can expand applications in treating deep-seated tumors and metastatic lesions.

While we did not evaluate toxicity in mice, in cell culture studies we clearly observed that the binding of FAP-α-IR700 to FAP-α expressing cells did not induce cytotoxicity. Cell death only occurred when these cells were exposed to light. Because we localized light exposure only to the tumor, cells outside of the tumor with FAP-α expression were not affected. This is different from approaches mentioned in the introduction where the FAP-α cytotoxic cargo is delivered systemically and targets all FAP-α expressing cells.

In conclusion, FAP-α-targeted NIR-PIT provides a novel and specific approach for eliminating FAP-α expressing CAFs in studies designed to understand the impact of these CAFs in tumor immune surveillance and progression. With increased availability of intra-operative or catheter-based light delivery and detection systems, the translation of this approach can provide a treatment strategy against a ubiquitous target to use in combination with immune checkpoint inhibitors for cancer treatment. The availability of FAP-α-PIT may also find uses in debilitating diseases such as fibrosis and arthritis.

## Materials and Methods

### Reagents

Water soluble phthalocyanine dye, IRDye 700DX NHS ester (IR700), was obtained from Li-Cor Bioscience (Lincoln, NE, USA). Anti-FAP-α polyclonal sheep antibody, AF3715, monoclonal mouse antibody, MAB3715, and monoclonal rat antibody, MAB9729 were purchased from R&D systems (Minneapolis, MN, USA). Anti-FAP-α monoclonal mouse antibody BMS168 was purchased from eBioscience (San Diego, CA, USA). Anti-FAP-α antibodies, ab137549, ab218164, ab28244, ab53066, ab207178, and ab227703 were purchased from abcam (Cambridge, MA, USA). Anti-FAP-α polyclonal rabbit antibody, PA5-51057 and sheep IgG isotype control (Cat. No. 31243) were purchased from Thermo Fisher Scientific (Waltham, MA, USA).

### Cloning and lentivirus production

The lentiviral vector pMA3211 was purchased from Addgene (Watertown, MA, USA). Cloning was outsourced to GenScript (Piscataway, NJ, USA). In brief, a PGK promoter was synthesized to replace the original TRE-Tight promoter via XhoI/SalI. Human FAP-α (Accession No. NM_004460) or murine FAP-α cDNA (Accession No. NM_007986.3) was synthesized and inserted into pMA3211 via SalI/NotI.

Lentivirus was produced and harvested according to our previously published method (Balaji Krishnamachary et al., 2009). Viral supernatants were derived by transient co-transfection of 293T (6×10^6^ in 100 mm^3^ petri-plates) cells using lipofectamine 2000 (Thermo Fisher Scientific). A total of 19.5 μg of plasmid in the proportion of 12 μg of lentiviral plasmid carrying human/murine FAP-α cDNA, 6 μg of packaging plasmid pCMVΔR8.2 and 1.5 μg of pCMV-VSVG was used, and viral supernatant collected at 48, 72 and 96 h post-transfection. Pooled supernatants were concentrated using an Amicon Ultra-15 (100 K cutoff) filter device (Millipore).

### Cell transduction and sorting

For lentiviral transduction, 2×10^6^ MDA-MB-231 or HT-1080 cells were plated onto 100 mm^3^ dishes and 5 ml of 10X concentrated viral supernatant with 1 mg/ml of polybrene was added for 4-5 h. This procedure was repeated for three days. Cells were maintained in culture medium containing 4 μg/ml of puromycin for selection. To sort out high FAP-α expressing cells, 4×10^7^ puromycin-selected cells were first incubated with 40 μg of AF3715, and then stained with phycoerythrin (PE)-conjugated anti-sheep secondary antibody IgG (F0126, R&D systems). Cell sorting was conducted on a BD FACSAria IIu cell sorter (Franklin Lakes, NJ, USA), and cells with the highest 90% of PE signal were collected. The sorted FAP-α overexpressing MDA-MB-231 and HT-1080 cells were denoted as 231-FAP, and HT-1080-FAP, respectively. NIH/3T3 fibroblasts were lentivirally transduced with murine FAP-α using the same protocol but without flow sorting. Murine FAP-α overexpressing fibroblasts were denoted as 3T3-FAP.

### Cells

MDA-MB-231 human breast cancer cells (notated here as 231 cells), HT-1080 human fibrosarcoma cells, and NIH/3T3 murine fibroblasts were purchased from American Type Culture Collection (ATCC, Manassas, VA, USA). Human mammary fibroblasts (HMFs) were kindly provided by Dr. Gary Luker, University of Michigan-Ann Arbor. Patient-derived prostate CAFs (PCAFs) were purchased from Asterand (Detroit, MI, USA). CAF35, a primary culture of stromal fibroblasts established from surgically resected pancreatic cancer tissue, was a generous gift from Drs. William Matsui and Asma Begum, Johns Hopkins University School of Medicine. FAP-α overexpressing human cancer cells (notated as 231-FAP or HT-1080 FAP) or murine NIH-3T3 fibroblasts (notated as 3T3-FAP) were lentivirally transduced using a lentiviral vector pMA3211 containing human or murine FAP-α cDNA, with a pGK promoter and a puromycin resistance gene as previously described (B. Krishnamachary et al., 2020).

All cells were cultured in DMEM medium supplemented with 10% FBS (Sigma, St. Louis, MO, USA). Cells were maintained at 37 °C in a humidified atmosphere containing 5% CO_2_.

### Synthesis of IR700-conjugated antibodies and concentration determination

The synthesis of IR700-conjugated antibodies was performed as previously described (Jin et al., 2016). Briefly, 1 mg of AF3715 was first dispersed in 1 ml of 1x PBS containing 159.2 μg of IR700 (81.6 nmol, 1 mM in DMSO). The mixture was maintained overnight at 4 °C, and then loaded onto Amicon Ultra-0.5 (10K cutoff) filter units (Millipore, Burlington, MA, USA) to remove the unbound IR700 molecules. The purified and concentrated conjugate (FAP-α-IR700) was sterilized by filtering through 0.2 μm membranes. Sheep IgG isotype control was similarly conjugated with IR700, and the conjugate was denoted as IgG-IR700. The concentration of antibody and dye/protein ratio was calculated by measuring the absorbance at 280 nm (ε_antibody_ = 210,000 M^−1^cm^−1^) and 689 nm (ε_IR700_ = 165,000 M^−1^cm^−1^). The correction factor of IR700 at 280 nm was 0.095.

### RNA isolation, cDNA synthesis, and quantitative reverse transcription polymerase chain reaction (qRT-PCR)

Total RNA was isolated from cells using QIAshredder and RNeasy mini kits (Qiagen, Hilden, Germany). cDNA was prepared from 1 μg of RNA using iScript cDNA synthesis kit (Bio-Rad, Hercules, CA, USA). cDNA samples were diluted 1:10 and real-time PCR was performed using IQ SYBR Green supermix and gene-specific primers in the iCycler real-time PCR detection system (Bio-Rad). The Hs-FAP-α primer was designed by using Primer3Plus, and the Mm-FAP-α primer using previously published data (Fan et al., 2016). The expression of target RNA relative to the house-keeping gene hypoxanthine phosphoribosyltransferase 1 (HPRT1) for human cells was calculated based on the threshold cycle (Ct) as R = 2^−Δ(ΔCt)^, where ΔCt = Ct_target_ −Ct _HPRT1_, Δ(ΔCt) = ΔCt_target_ − ΔCt _wild type_. For mouse cells, fold expression was calculated relative to 18s RNA expression.

### Flow cytometry

Cells were detached using TrypLE (Thermo Fisher Scientific). Freshly resected tumor tissue was dissociated into single cell suspensions using a tumor dissociation kit (130-096-730, Miltenyi Biotec, Auburn, CA, USA) according to the manufacturer’s protocol. Cells were then dispersed at 1×10^6^ per 100 μl of FACS buffer made with 1x PBS supplemented with 1% BSA and 2 mM EDTA. For FAP-α staining, cells were incubated on ice for 30 min with 1 μg of AF3715. Polyclonal sheep IgG was used as control. After a single wash, cells were re-suspended in 100 μl of FACS buffer and incubated with 10 μl of PE-conjugated anti-sheep IgG secondary antibody (F0126, R&D systems) for 30 min on ice. For tumor-dissociated cells, prior to incubation with primary antibody, rat anti-mouse CD16/32 antibody (clone 2.4G2, BD Pharmingen™, San Diego, CA, USA) was added for Fc blocking. LIVE/DEAD™ Fixable Dead Cell Stain Kit (Thermo Fisher Scientific) was used after incubation with secondary antibody to identify and distinguish live cells from dead cells. Flow cytometry was conducted on a FACS Calibur (BD Bioscience, Franklin Lakes, NJ, USA) and analyzed by FlowJo software (FLOWJO, Ashland, OR, USA).

### Co-culture study

HMFs at a density of 8×10^4^/3 ml were plated in each well of a 6-well companion plate (Corning, Corning, NY, USA), and MDA-MB-231 cells at a density of 8×10^4^ cells/2 ml were seeded in each Falcon™ cell culture insert containing a 0.4 μm transparent polyester (PET) membrane (Corning). After a 3-day incubation, the HMFs were detached, re-seeded at a density of 8×10^4^/3 ml and co-cultured for a further 3 days with fresh MDA-MB-231 cells at a density of 8×10^4^ cells/2 ml. Following the second 3-day incubation, HMFs were collected for FAP-α and α-SMA immunostaining. FAP-α immunostaining was performed following the procedure detailed earlier for flow cytometry except for using Per-Cp conjugated anti-sheep IgG as secondary antibody. For α-SMA staining, HMFs were first fixed with 4% PFA and permeabilized by 0.4% Triton X-100. Immunostaining was performed with anti-α-SMA polyclonal rabbit antibody (ab5694, abcam) or rabbit IgG isotype control followed by staining with APC-conjugated anti-rabbit IgG secondary antibody (F0111, R&D systems).

### Confocal microscopy

Wild-type and FAP-α overexpressing cells were seeded in an 8-well Lab-Tek II chamber slide (Nalge Nunc, Rochester, NY, USA) at a density of 10,000 cells/well overnight, and incubated with FAP-α-IR700 or IgG-IR700 at a concentration of 5 μg/ml for 1 h at 37 °C. To investigate competition binding, a five-fold excess of AF3715 (5 μg/ml) was added to a separate set of wells 15 min prior to adding FAP-α-IR700. After a single wash, cells were fixed with 4% PFA and imaged with a laser scanning confocal microscope (Zeiss LSM 510-Meta, Carl Zeiss Microscopy GmbH, Jena, Germany). The red laser at 633 nm was used to excite IR700, and the receiving PMT channel was set at 680~700 nm. The IR700 fluorescence was displayed in pseudo magenta color. All the images were obtained under identical microscope settings.

### Immunoblot assay

Cells or homogenized tumor tissue were lysed in radioimmune precipitation (RIPA, Sigma) buffer and measured by a BCA assay (Pierce) for protein concentration. Cell lysate at 100 μg of protein in 1x loading buffer with β-mercapthoethanol was boiled for 50 min at 95 °C. Denatured protein was later resolved by SDS-PAGE and transferred to a nitrocellulose membrane. A recombinant anti-FAP-α monoclonal rabbit antibody ab207178 (clone EPR20021, abcam) was used to probe human/murine FAP-α. GAPDH or Actin was used as loading control.

### Cell viability

The specificity and effectiveness of FAP-α-IR700-PIT were evaluated using cell viability assays. In a typical assay, cells were seeded overnight in 96-well plates at a density of five thousand cells/well. Cells were further incubated for 1 h at 37 °C in medium containing either FAP-α-IR700 or IgG-IR700 or AF3715. After carefully aspirating the medium and replenishing with fresh media, cells were exposed to light using a light emitting diode (LED, Marubeni, Tokyo, Japan) that provided continuous NIR irradiation at 690 nm. The power of the light exposure was measured by an optical power meter (PM 100, Thorlabs, Newton, NJ, USA). Immediately after NIR light exposure, 10 μl of CCK-8 reagent (Dojindo, Mashiki, Japan) was added to each well for 3 h and the absorbance at 450 nm was measured on an Epoch™ Microplate Spectrophotometer (Biotek, Winooski, VT, USA). Cytotoxicity data were expressed as mean ± standard derivation (SD) from at least triplicate wells. In studies characterizing antibody concentration or light-dose dependency, the concentration of FAP-α-IR700 was varied from 0.1 to 5 μg/ml or light intensity was varied from 2 to 8 J/cm^2^. The specificity of FAP-α-IR700-mediated phototoxicity in comparison with wild-type MDA-MB-231 and HT-1080 cells, or unconjugated AF3715 antibody or IgG-IR700 was established. In a separate study, plates with cells were wrapped in aluminum foil to evaluate effects without light exposure. The effect of competitive inhibition was also examined by adding 5x AF3715 15 minutes prior to FAP-α-IR700.

### Tumor models

Animal studies were conducted in accordance with approved protocols. Six-to eight-week-old female athymic Balb/c (nu/nu) mice were purchased from Charles River (Wilmington, MA, USA). Tumor xenografts from the cancer cell lines were established bilaterally by inoculating 1×10^6^ cancer cells in 0.1 ml of Hanks balanced salt solution in the second mammary fat pad for 231 and 231-FAP tumors or in the flank for HT-1080 and HT-1080-FAP tumors.

### *In vivo, ex vivo* fluorescence imaging and biodistribution

NIR fluorescence imaging was performed on a Li-Cor Pearl® Impulse imager (LI-COR Biosciences). Mice bearing bilateral 231 and 231-FAP tumors (n = 4 per group) or HT-1080 and HT-1080-FAP tumors (n = 3 per group), were imaged once tumor volume reached 100 mm^3^. Next, 50 μg of FAP-α-IR700 or IgG-IR700 was injected intravenously (*i*.*v*.) through the tail vein, and fluorescence images were obtained over a 24 h period at 0, 1 h, 6 h, 24 h post-injection (*p*.*i*.). At 24 h *p*.*i*., mice were sacrificed, and major organs and tumors were resected for *ex vivo* imaging. Images were acquired under identical experimental conditions. Regions of interest were drawn on *ex vivo* images and analyzed by Pearl Impulse software (Li-Cor Biosciences) to determine fluorescence intensity. Bio-distribution values were normalized to the weight as % injected dose/g (%ID/g) from three or four mice per group (n = 3 or 4), using a calibration curve of intensity *versus* blood FAP-α-IR700 concentration.

### *In vivo* PIT

Once tumor volumes reached approximately 100 mm^3^, tumor-bearing mice were randomly assigned to three groups (n = 4 per group) based on the different injections: (i) PBS; (ii) FAP-α-IR700; (iii) IgG-IR700. Next, 100 μg of antibody conjugate or 100 μl of PBS were injected *i*.*v*. into each mouse on day 0 and again on day 7. NIR light exposure at a power of 200 J/cm^2^ was given at 24 h *p*.*i*. Caliper measurements of tumor volumes were obtained over a 2-week period on Day 0, 3, 7, 10, and 14, following which mice were euthanized and the tumors excised for immunohistochemistry (IHC), hematoxylin and eosin (H&E) staining, and western blot analysis.

### Immunohistochemistry

Human breast cancer tissue microarrays (TMAs, BR243k) from 6 cases of breast invasive ductal carcinoma (two cores per group) with matched adjacent breast tissue (two cores per group) were purchased from US Biomax (Derwood, MD, USA). The TNM stage and grade of 6 cases were TisN0M0 and grade 1 for cores A1 and A2, T2N2M0 and grade 2 for cores A5 and A6, T3N0M0 and grade 2 for cores B1 and B2, T2N2M0 and grade 3 for cores B5 and B6, T2N0M0 and grade 2 for cores C1 and C2, T2N0M0 and grade 3 for cores C5 and C6. TMA slides were immunostained for FAP-α according to standard IHC protocols.

Five-micron tumor sections obtained from formalin fixed paraffin embedded xenografts were stained with H&E, and for FAP-α according to standard IHC protocols. Antigen retrieval was performed by boiling the slides in citric buffer at pH 6 for 50 minutes. Anti-FAP-α antibody, ab207178 (abcam, 1:300 dilution) was used for immunostaining both human and murine FAP-α. Slides were digitally scanned at 20X magnification and analyzed by Aperio ImageScope software (Leica Biosystems, Richmond, IL, USA).

### Statistical analysis

Data were expressed as mean ± SD from three or more samples or three or more mice. Statistical analysis was performed with a two-sided student t-test (Microsoft Excel, Redmond, WA, USA), assuming unequal variance. Values of *P* ≤ 0.05 were considered significant, unless otherwise stated.

## Supporting information

**Supplementary files:** Supplementary Table 1-Source Data Files.

## Acknowledgements

We thank Drs. William Matsui and Asma Begum for generously providing CAF35 cells. We thank Mr. G. Cromwell for his valuable technical assistance. We thank Dr. K. M. Horton for her support.

## Notes

**Financial Support:** This work was supported by NIH R35 CA209960, R01 CA82337, P41 EB024495, P30 CA006973, ZIA BC011513 and a grant from the Emerson Collective Cancer Research Fund.

**Conflict of Interest Disclosures:** The authors have no conflict of interest to disclose

### Competing Interest Statement

The authors have declared no competing interest.

